# Assessment of environmental impacts based on particulate matter, and chlorophyll content of urban trees

**DOI:** 10.1101/2023.05.31.543026

**Authors:** Vanda Éva Abriha-Molnár, Szilárd Szabó, Tibor Magura, Béla Tóthmérész, Dávid Abriha, Bianka Sipos, Edina Simon

## Abstract

Trees improve air quality, and they have irreplaceable aesthetic value in urban landscapes. The amount of dust deposited on tree leaves is a simple and cost-effective indicator of air quality. Our aim was to explore particle filtering capacity of roadside trees in an urbanized area. We also assessed the impact of meteorological conditions on the amount of deposited dust. We measured the leaf surface deposition, and chlorophyll content of leaves along a road section that started at an intersection and ended in a less disturbed suburban area in Debrecen city, Hungary. Samples were collected in July, and September from *Celtis occidentalis*, a frequent species. We found a significant negative correlation between dust deposition on the leaves, and the distance from the intersection in July, meaning that the amount of dust on the leaves decreased as the distance from traffic increased. In September, dust deposition decreased considerably compared to July, caused by the rainfall before the second sampling. Chlorophyll content also had a significant negative correlation with the distance from the traffic intersection in July, as it decreased towards the less disturbed end of the transect. We also found a positive correlation between dust deposition and chlorophyll content in July. Surprisingly, the exposure to moderate amounts of pollutants in the air caused an increase in chlorophyll content. Our findings suggest that dust deposition on leaves serves as a reliable indicator of traffic intensity, because excess dust caused by the proximity of car traffic can be detected on the leaf surface. Although, certain weather conditions like rainfall and wind can disrupt the patterns in dust deposition that have developed over an extended period through wash-off and resuspension. Hence, it is advisable to consider these effects while selecting the sampling time and evaluating the results.

**Highlights:**

- Leaves of urban trees are used as bioindicators of deposited dust pollution.
- Dust deposition, and chlorophyll content was measured along a roadside transect.
- We found that dust and chlorophyll content decreased with distance from traffic emissions.
- Moderate level of dust pollution resulted in an increase in chlorophyll content.

## 1. Introduction

One of the most remarkable air pollutants in large cities is particulate matter. Particulate matter with aerodynamic diameters between 0.001 and 100 µm can occur in air (Klejnowski et al., 2013). Usually, in its settled state, it is collectively referred to as deposited dust, regardless of size. A wide range of inorganic and organic contaminants are likely to be found in the deposited dust (Alghamdi et al., 2023). The main anthropogenic sources include traffic, domestic heating, various construction and renovation projects, and agricultural activities in the areas surrounding the city. Fine particles are mainly originated from combustion, while coarser particles are more likely to be the result of mechanical abrasion and resuspension (Braun et al., 2007).

In major cities concentrations of common pollutants are often monitored at fixed point stations (Nagendra et al., 2021). The EU air quality standard for the annual PM10 concentration is 40 µg/m^3^. However, even if the current and/or average concentrations of pollutants are below the respective thresholds, the synergistic effect between them can be more harmful to living organisms than the individual pollutants. Moreover, the number of monitoring stations is often insufficient; thus, they may not representative of the municipality as a whole (Kumar et al., 2015).

Green spaces can effectively reduce air pollutants (Lei et al., 2018). This is one of the ecosystem services that can be directly linked to human health impacts and can even have an associated monetary value (Fusaro et al., 2017; Nowak et al., 2014; Sebastiani et al., 2021). Air pollutants can negatively affect the growth, pigmentation, and photosynthetic activity of the plant as anthropogenic stressors in high concentrations (Bharti et al., 2018). Urban trees tend to be easily damaged, and above certain concentrations and exposure times, visible alterations such as chlorosis, reduction in leaf number, leaf area, stem and root length can occur (Joshi and Swami, 2007). Chlorophyll is a vital pigment for photosynthetic activities and its content is usually influenced by certain air pollutants, depending on the species (Sen et al., 2017). Therefore, in sensitive species, it is a commonly used indicator of air quality.

Biomonitoring is an accessible low-cost solution for air quality assessment, although it has the limitation that a survey only provides information on the pollution status at the time of sampling interpreted for the prior vegetation period. *Celtis occidentalis* L. has been used as a bioindicator in the past to assess environmental pollution in urban areas (Greksa et al., 2019; Molnár et al., 2020; Simon et al., 2014). Its wide spatial distribution makes it an appropriate candidate for biomonitoring. Simon et al. (2014) also showed that this species had a high density of trichome on the leaf surface, which resulted in a high dust absorbing capability. Other morphological traits, such as stomata density and roughness, contribute to the accumulation of particulate matter (Sgrigna et al., 2020).

Dust deposition on urban vegetation is definitely in the focus of researchers worldwide. Research objectives can generally be categorized into two groups: biomonitoring and pollution reduction, which are closely related and overlap to some extent. The dust deposition on roadside tree canopies has already been investigated, for instance, to detect barrier function (Tong et al., 2016; Zheng et al., 2021), and to mitigate, or biomonitor traffic emissions (Bui et al., 2023; Ozdemir, 2019). Furthermore, the dilution of particulate matter with distance from high traffic roads has also been studied, mostly using monitoring stations (Karner et al., 2010; Patton et al., 2014).

Our objective was to assess how effectively a row of trees at different distances from an intersection filtered particulate pollution. Intersections generally produce increased levels of pollutants due to the constant acceleration and deceleration of vehicles. The aim of our study was to analyze leaf surface deposition and chlorophyll content of roadside trees in a unique layout in Debrecen. Trees planted along a less frequented street section were sampled, where one end of this transect was connected to a high-traffic, signalized intersection, while the other end led to a suburban area. We aimed to find out whether the excess dust generated by the vehicles could be detected on tree leaves in the presence of other urban impacts. Samples were collected once during a dry, rain-free period in July and once after rainfall in September to assess the impact of meteorological factors. Our hypotheses were: (i) the amount of dust deposition is higher on leaves collected near the intersection compared to the suburban area, (ii) after rainfall, the dust deposition is reduced compared to the dry period, (iii) chlorophyll content is higher in leaves collected in the suburban area compared to the intersection, and (iv) there is a positive correlation between dust deposition and chlorophyll content.

## 2. Materials and methods

### 2.1. Study area and sample collection

The study site was in the central part of Debrecen city. It is at 120 m above sea level on the Great Hungarian Plain (Lóki, 2020). Debrecen has limited area of urban green spaces that are scattered throughout the city (Csomós et al., 2020). Currently there is no significant industrial pollution source in the region, but several major industrial investments are planned in the near future. The air quality in the city is mostly affected by the particulate pollution from vehicular traffic. Nearby agricultural activities also have a large contribution to pollution, especially from the western side, where there is no vegetation functioning as a windbreak. The city is also neighboring to the Nyírség region, which is characterized by high erodibility by wind, and therefore contributes an additional source of particulate matter (Négyesi et al., 2019). Debrecen’s air exchanges slowly and is not easily renewed due to geographical and meteorological conditions and general urban effects. Also, a large amount of mineral dust from the Sahara reaches Hungary every year (Varga, 2020). Nevertheless, the annual average concentration of continuously monitored particulate matter in the city has not exceeded the health limit value in recent years.

Sampling was carried out along a 400 m section of a street, with a starting point approximately 100 m from a busy intersection, and an end point in a less disturbed suburban area, in July and September 2022 (Figure 1). The Hungarian Meteorological Service reported 16.5 mm of precipitation during the 5 days prior to sampling in September (OMSZ 2023). Leaf samples were taken every 50 m along the chosen road section in three replicates from common hackberry trees (*Celtis occidentalis* L.) which is a deciduous tree planted along roadsides. There was a total of 54 samples; each sample contained approximately 10 leaves. The samples were stored in paper bags and kept frozen until further processing.

**Figure 1.**
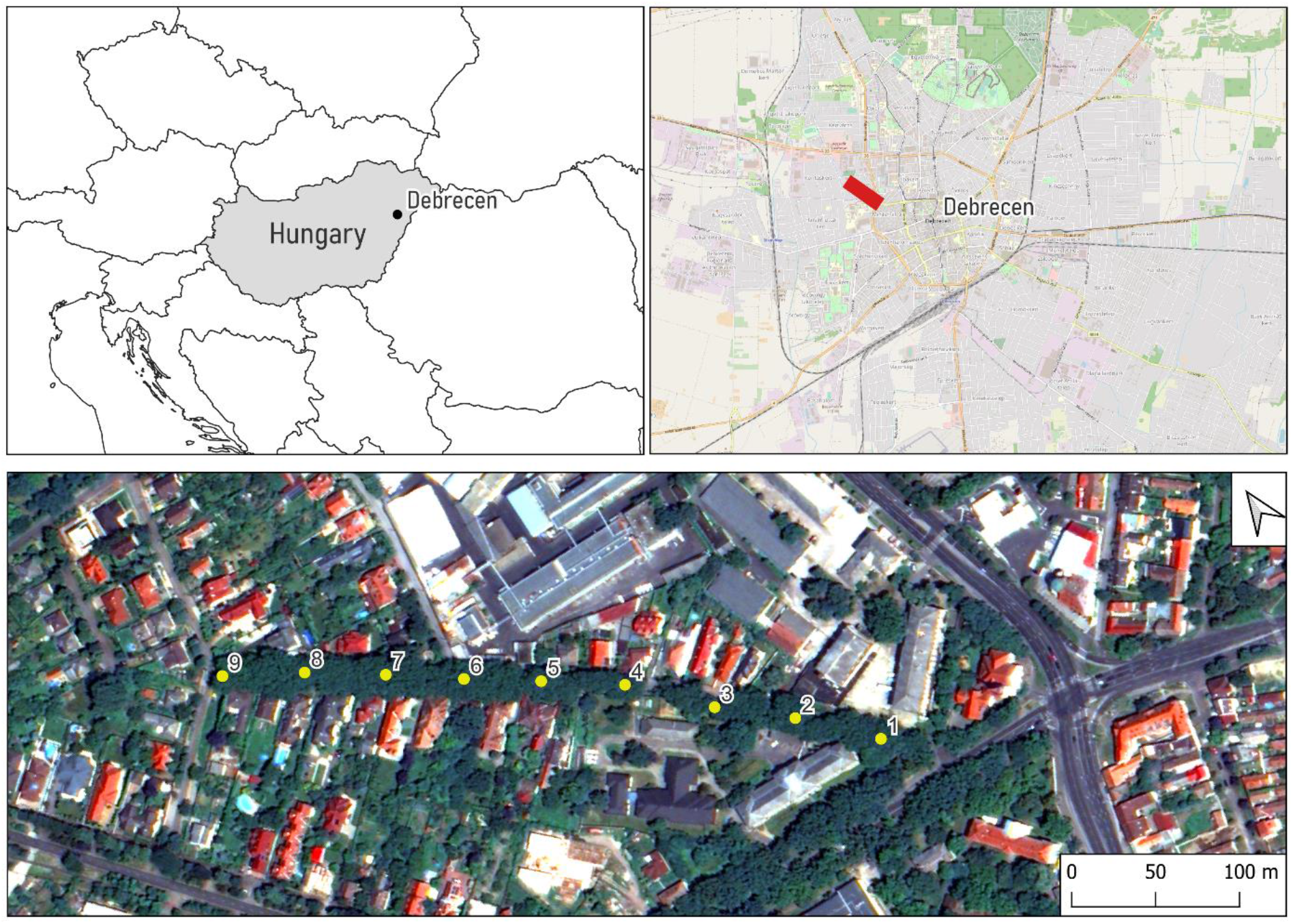
The location of sampling sites along the transect in Debrecen, Hungary.

### 2.2. Analysis of dust deposition

Leaf area of each leaf was determined. Then, leaf samples were placed in 500 ml plastic containers, and mixed for 10 minutes on an analog orbital shaker (GFL 3015) with 250 ml deionized water. The resulting suspension was filtered through a 100 μm sieve and the procedure was repeated with another 50 ml of deionized water (Simon et al., 2011, 2014). The resulting 300 ml suspension was then filtered through a vacuum pump (BOECO R-300) using a filter paper of 5-8 µm retention diameter (Munktell 392, Ahlstrom), previously weighed on an analytical balance (ME, METTLER TOLEDO). The filter papers were reweighed with the collected dust to obtain the net mass of dust, which was converted into μg/cm^2^ of leaf area.

### 2.3. Analysis of chlorophyll content

For the analysis of chlorophyll content, approximately 20 mg of fresh leaf tissue was prepared from each sample. The leaf tissue was crushed in a mortar and homogenized in 5 ml of 96% (v/v) ethanol. The extracts were centrifuged at 1500 rpms for 3 minutes (IEC Centra MP4). The absorbances were measured using spectrophotometric analysis (BOECO S-220) on the wavelengths of 653, 666, and 750 nm against 96% (v/v) ethanol blank. Chlorophyll content was calculated based on Equation 1, where *V* is the extract volume, *m* is the fresh weight of leaf tissue, and *E666* and *E653* are the absorbances at 666 nm and 653 nm minus the absorbance at 750 nm, respectively.

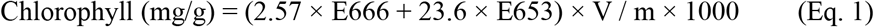

### 2.4. Statistical analysis

We used SPSS Statistics 21 (IBM Corp., Armonk, NY, USA) software for statistical analyses. Normal distribution of the variables was checked by the Shapiro-Wilk test. Pearson’s correlation coefficient was used to reveal the relationship among variables (distance from sampling site, dust deposition and chlorophyll content). Simple regression analysis was used to describe the relations between the variables. When error variances showed heteroscedasticity weighted least squares regression was used, where the weights were based on the reciprocal of the given variable’s variance. The difference between samples in July and September was tested using the paired t-tests.

### 3. Results

Based on the samples from July, average dust deposition ranged widely from 4.5 ± 1.7 µg/cm^2^ to 99.6 ± 41.2 µg/cm^2^ along the sampled transect (*Figure 2a*). There was a significant negative correlation (r = –0.847, p < 0.001) between the dust deposition on the leaves and the distance from the intersection. We found the lowest average dust deposition at the sampling point furthest from the intersection. The trend of change was not completely monotonically linear, as e.g., the second closest point to the intersection had the highest average dust deposition. Based on the weighted regression analysis, 80.8% of the variance in dust deposition could be explained by the model (p < 0.001). The regression model predicted an average decrease of 10.5 µg/cm^2^ in dust deposition every 50 m from the intersection, which was the approximate distance between each consecutively sampled tree.

**Figure 2.**
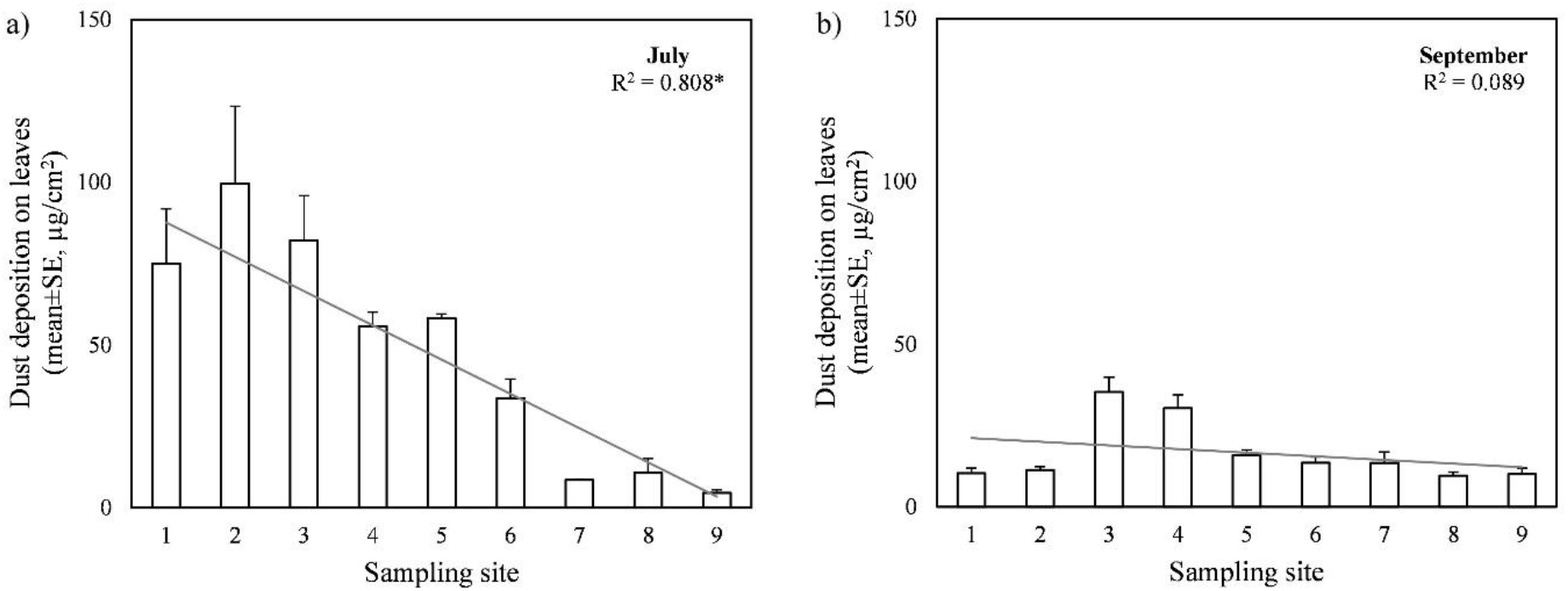
Dust deposition on leaves collected a) in July and b) in September, with the fitted regression line and the goodness-of-fit value (R^2^). Asterisks (*) indicate statistically significant (p < 0.05) relationship. The sampling sites were 50 m apart and the 1^st^ one was closest to the intersection.

Dust deposition on the leaf surface differed significantly between the sampling dates in July and September (p < 0.001). In September, dust deposition decreased by 30.9 µg/cm^2^ on average compared to July (*Figure 2b*). We observed the largest absolute differences at the sampling points closest to the intersection where the dust deposition was initially high in July. In this case, the correlation between dust deposition on the leaves and distance from the intersection was not significant, due to the variance and the non-linear trend of the relationship. Thus, our first hypothesis about the dust exposure near the intersection was only confirmed by the first assessment in July. Our hypothesis about the wash-off effect of precipitation is also confirmed.

Similar to dust deposition, chlorophyll content also had a significant negative correlation (r = –0.813, p < 0.001) with the distance from the traffic intersection in July. At this time of sampling, chlorophyll content ranged from 3.03 ± 0.67 mg/g to 7.37 ± 1.21 mg/g in the leaves of *C. occidentalis* (at the sampling points furthest and nearest to the intersection, respectively). In this case, 70.2% of the variance in chlorophyll content could be explained by the weighted regression model (p < 0.001) (*Figure 3a*). The model predicted an average decrease of 0.51 mg/g in chlorophyll content every 50 m from the intersection. Chlorophyll content also differed significantly between the sampling dates (p < 0.001). In September, chlorophyll content was higher than in July by 2.0 mg/g on average across the sampling points (*Figure 3b*). There was a considerable variability among the sampled trees, and this time, correlation with the distance from traffic was not significant. Based on these results, we rejected our hypothesis that chlorophyll content is higher in the less disturbed area.

**Figure 3.**
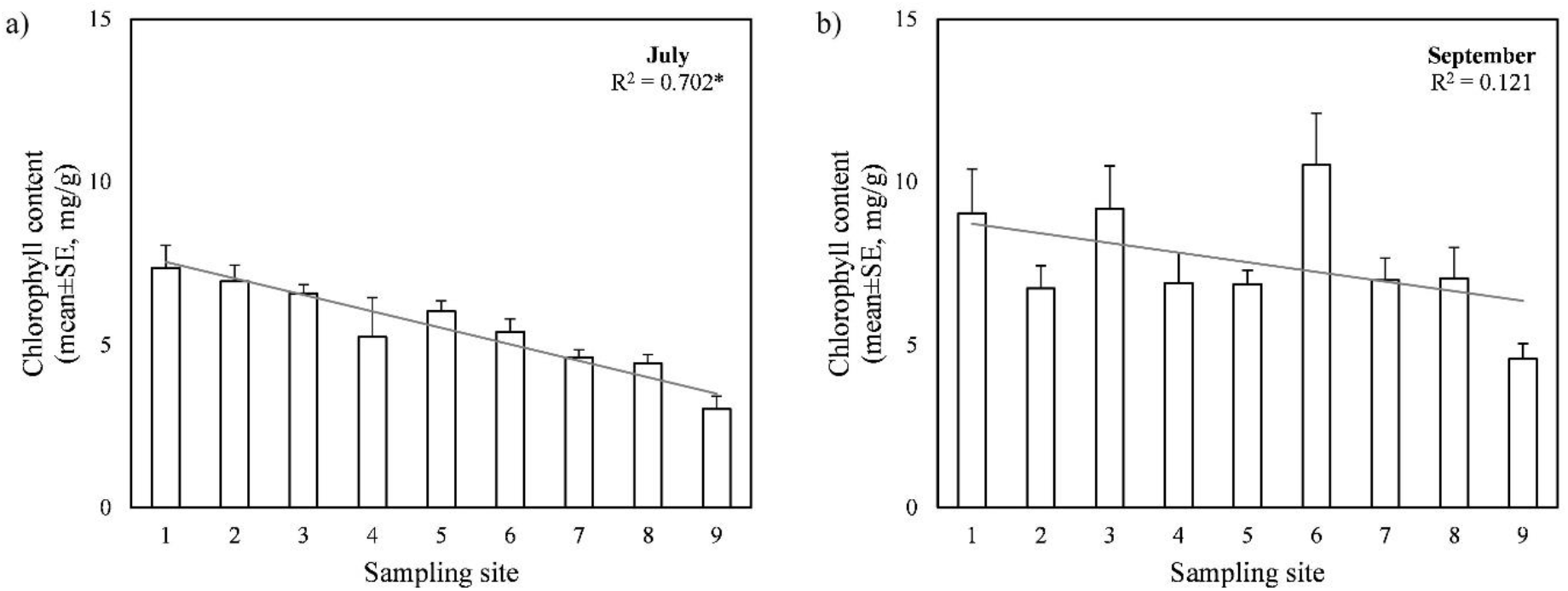
Chlorophyll content of leaves collected a) in July and b) in September, with the fitted regression line and the goodness-of-fit value (R^2^). Asterisks (*) indicate statistically significant (p < 0.05) relationship. The sampling sites were 50 m apart and the 1^st^ one was closest to the intersection.

Lastly, we explored the relationship between dust deposition and chlorophyll content from the same samples for the entire transect. Based on the samples collected in July, we observed a significant positive correlation (r = 0.684, p < 0.001) between the dust deposition and the chlorophyll content of the leaves, which means that as the mass of deposited dust on the leaf surface increased, the concentration of chlorophyll increased in the leaf tissue as well. Regression analysis showed a significant, but relatively weak relationship, with R^2^ = 0.468 (p < 0.001) (*Figure 4*). In September, there was no significant correlation between dust deposition and chlorophyll content.

**Figure 4.**
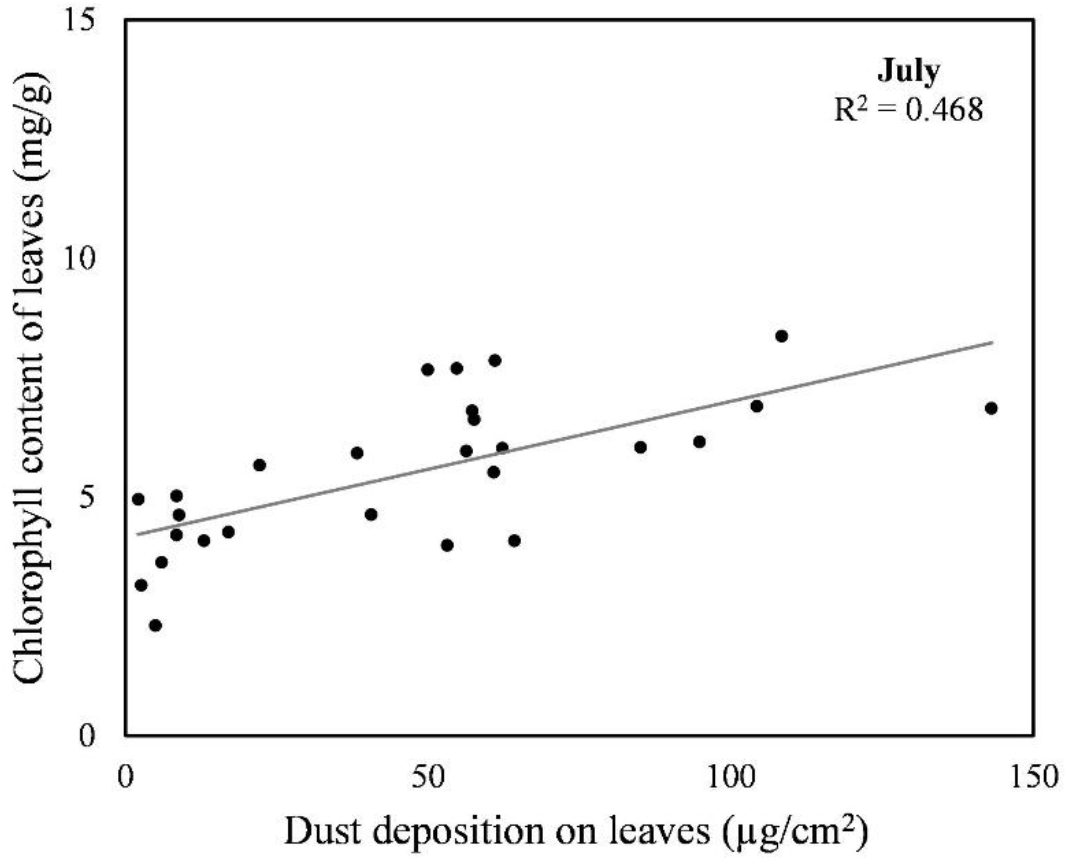
Regression plot for the dust deposition and the chlorophyll content of leaves collected in July.

## 4. Discussion

### 4.1 Dust deposition on roadside tree leaves

We assessed the effects of vehicular traffic on air quality by determining the dust deposition accumulated on leaves of roadside trees. We found that the dust deposition on *C. occidentalis* leaves decreased the further it was sampled from the intersection, which indicated continuous dilution and deposition of traffic-generated and resuspended dust with distance. The pollution source in our study was the traffic flow through intersection near the beginning of the sampling. The road where the samples were collected had sparse traffic; thus, its contribution to dust emissions was negligible, leaving the traffic flow through intersection as the main source of dust emission. In a meta-analysis, Cai et al. (2017) concluded that leaf deposition negatively correlated with distance from roadsides or any pollution source and our result was consistent with the above statement. The concentration of solid particles naturally dilutes with distance from the road due to dispersion, but the presence of vegetation promotes this process by filtering and collecting particles. It is important to mention that to avoid large elevations in on-road concentrations, roadside vegetation should not be so dense as to form a solid barrier for particles (Tong et al., 2016; Zheng et al., 2021).

Our results in September showed a general decrease in dust deposition compared to July. The weather before the second sampling turned windy and rainy, which presumable led to wash-off and caused this reduction. Based on data from the Hungarian Meteorological Service, in 5 days prior to the sampling in July there was 0.7 mm of precipitation, while before the sampling in September there was 16.5 mm (OMSZ 2023). However, the rate of change was not consistent at the sampling points, as the largest decrease was observed at points closer to traffic, because that end of the road is more exposed to meteorological conditions and turbulent winds caused by traffic. At both sampling times, the maximum average dust deposition was observed only at the 2nd or 3rd point from the intersection. This indicates that the dust stirred up by traffic is transported in the air for a certain distance and only then settles on the leaf surface. There are also several other environmental factors that influence the amount of dust deposition. For example, samples were collected from the same height, but the different crown size and overall leaf surface above the sampling level had varying effect on the deposition and the transfer of dust particles during a raining event. The effect of rain is an important factor to consider when discussing dust retention by plants. The accumulation of dust particles saturates the leaf surface over time, but the amount of dust will decrease again due to rainfall and wind. The exact measurement of this exchange is limited, and it is generally carried out using simulated rainfall in an artificial environment. For example, Weerakkody et al. (2018) found that particles were washed off of leaves during simulated rainfall. However, Wang et al. (2015) determined that natural rainfall can also increase the accumulation of particles on the leaf surface at high concentration of PM pollution. This is usually the case during low amount of rainfall, when the water-holding capacity of the leaf is not exceeded (Cai et al., 2017). In contrast, Freer-Smith et al. (2015) even suggested that particles were not easily removed by rain once deposited on the leaf surface. It can be assumed that the effect of rain wash-off will not always be obvious, and that different external circumstances can change the amount of dust deposition in distinctly different directions.

### 4.2 Chlorophyll content of tree leaves

Along with the dust deposition, chlorophyll content was also determined from the same samples along the road. We found that chlorophyll content of *C. occidentalis* decreased with the distance from the traffic intersection in July; although, at a lower rate than the dust deposition did. This indicates that exposure to moderate amounts of pollutants in the air has triggered an increase in chlorophyll content. This is not necessarily an unusual finding, although several studies have reported the contrary. Air pollutants are often found to cause a depletion of chlorophyll in various plant species (Chaudhary and Rathore, 2019; Giri et al., 2013; Sen et al., 2017). However, similarly to our results, Petrova et al. (2011) found that chlorophyll in *Betula pendula* leaves was present in higher concentrations in areas with more intense anthropogenic activity. It was suggested that below a certain level of air contaminants, chlorophyll content is positively affected due to compensatory mechanisms. Paull et al. (2021) found similar chlorophyll content in green wall plants at test and control sites. This was explained by the innate tolerance of plants and the relatively good air quality, which is also true for the present study area.

Compared to July, chlorophyll content of *C. occidentalis* slightly, but significantly increased in September. However, the quantities determined at each location showed a high degree of irregularity in relation to each other, thus, unlike in July, no definite trend was discernible along the sampling transect in September. Chlorophyll content in the leaves of deciduous trees changes naturally with the seasons. In the local climate, Szőllősi et al. (2010) found that in the leaves of *Quercus petraea* (Matt.) Liebl., chlorophyll content only slightly fluctuated after the initial increase in the beginning of the growing season until September. However, in urban environments, anthropogenic influences modify the natural variation of chlorophyll to some extent. For example, You et al. (2016) found that while chlorophyll concentration in *Platanus occidentalis* L. increased slightly from June to October at an unpolluted site, it decreased at a polluted site in that same time period.

### 4.3 The relationship between dust deposition and chlorophyll content

The relationship between the dust deposition and the chlorophyll content can be approached from different directions. In July, the chlorophyll content has varied in parallel with the dust deposition on the leaf surface, while there was no clear relationship between the two parameters in September. This behavior is usually dependent on the tree species. On the one hand, some species show a decrease in chlorophyll in the presence of pollutants, while others increase chlorophyll as a defense mechanism to maintain photosynthetic activity. *C. occidentalis* is well known for its resilience in urban environments, which explains this latter behavior. On the other hand, in terms of temporal variations, an increase in chlorophyll content was observed along with a decrease in dust deposition on the leaf surface over the time of the observations. Most literature on this subject comes from India where air quality is frequently poor (Banerjee et al., 2021). Chlorophyll content of leaves is reported to be lower at roadsides and industrial areas with higher levels of pollution compared to control areas (Chaurasia et al., 2013; Gupta et al., 2016; Hariram et al., 2018; Rai and Panda, 2014). However, dust deposition in most of the studies are in the magnitude of a few mg/cm^2^, while even the highest amount we found was only 99.6 ± 41.2 µg/cm^2^. Therefore, it is assumed that the air quality and dust pollution at the sampling site in Debrecen was not severe enough to cause a deficit in the chlorophyll content of leaf tissue.

## 5. Conclusions

Our models predicted an average decrease of 10.5 µg/cm^2^ in dust deposition every 50 m from the intersection. The dust deposition on the surface of tree leaves is a good reflection of the degree of dust pollution generated by traffic. However, it is very important to take meteorological conditions into account in such studies. In many cases, information on precipitation and wind conditions is missing from the description of the sampling, which compromises the comparability of the results with others. This is also demonstrated in the present study, as the first sampling yielded the expected result for dust exposure, while the later date did not lead to a clear correlation. In addition, further research on the relationship between chlorophyll content and air pollution is still necessary, as different geographic factors, specific characteristics of species and varying levels of pollution in an area have a strong influence on the alteration in chlorophyll content. Therefore, any conclusion about air quality based on chlorophyll content needs careful consideration, as chlorophyll levels can increase with low levels of pollution.

Overall, it was concluded that dust deposition on the leaves is a good indicator of the intensity and proximity of traffic, although certain weather conditions (rainfall, wind) can disrupt patterns that have developed over a longer period of time. It is therefore recommended to consider these effects when choosing the sampling time and when evaluating the results.

## Author contributions

**V.É.A-M**.: Conceptualization, Formal analysis, Writing - original draft. **S.S**.: Writing - Review & Editing. **T.M**.: Methodology, Writing - Review & Editing. **B.T**.: Writing - Review & Editing. **D.A**.: Formal analysis, Writing - Review & Editing. **B.S**.: Investigation, Writing - Review & Editing. **E.S**.: Resources, Supervision, Writing - Review & Editing.

## Acknowledgements

The research is supported by the ÚNKP-22-4-II New National Excellence Program of the Ministry for Culture and Innovation from the source of the National Research, Development and Innovation Fund. SS and DA was supported by the NKFI K 14121 project.

